# SARS-CoV-2 Envelope protein (E) binds and activates TLR2: A novel target for COVID-19 interventions

**DOI:** 10.1101/2021.11.10.468173

**Authors:** Rémi Planès, Jean-Baptiste Bert, Sofiane Tairi, Lbachir Benmohamed, Elmostafa Bahraoui

**Affiliations:** INSERM, U1043, CPTP, CHU purpan, Toulouse, France; CNRS, U5282 CPTP, CHU purpan, Toulouse, France; Université Paul Sabatier, CPTP, CHU purpan, Toulouse, France; IPBS, Toulouse, France; Laboratory of Cellular and Molecular Immunology, Gavin Herbert Eye Institute, University of California Irvine, School of Medicine, Irvine, CA, 92697, United States of America

## Abstract

In this study, we present a molecular characterization of the interaction between the SARS-CoV-2 envelope protein E with TLR2. We demonstrated that E protein interacts physically with TLR2 receptor in a specific and dose-dependent manner. Furthermore, we showed that this interaction is able to engage TLR2 pathway as demonstrated by its capacity to activate NF-κB transcription factor and to stimulate the production of CXCL8 inflammatory chemokine in a TLR2-dependent manner. Furthermore, in agreement with the importance of NF-κB in TLR signaling pathway, we showed that the chemical inhibition of this transcription factor led to significant inhibition of CXCL8 production, while blockade of P38 and ERK1/2 MAP kinases resulted only in a partial CXCL8 inhibition. Overall, our findings suggest considering the envelope protein E as a novel target for COVID-19 interventions: (*i*) either by exploring the therapeutic effect of anti-E blocking/neutralizing antibodies in symptomatic COVID-19 patients, or (*ii*) as a promising non-Spike SARS-CoV-2 antigen candidate to include in the development of next generation prophylactic vaccines against COVID-19 infection and disease.

**Importance:** Although, the exact mechanisms of COVID-19 pathogenesis are unknown, recent data demonstrated that elevated levels of pro-inflammatory cytokines in serum is associated with enhanced disease pathogenesis and mortality. Thus, determining the molecular mechanisms responsible for inflammatory cytokine production in the course of SARS-CoV-2 infection could provide future therapeutic targets. In this context, to the best of our knowledge, our report is first to use a detailed molecular characterization to demonstrate that SARS-CoV-2 Envelope E protein binds to TLR2 receptor. Specifically, we showed that SARS-CoV-2 Envelope E protein binds to TLR2 in a direct, specific and dose-dependent manner. Investigating signalling events that control downstream activation of cytokine production show that E protein / TLR2 binding leads to the activation of NF-κB transcription factor that control the expression of multiple pro-inflammatory cytokines including CXCL8. Overall, our findings suggest considering the envelope protein E as a novel target for COVID-19 interventions.

## 1. Introduction

SARS-CoV-2, the etiologic agent of the current worldwide COVID-19 pandemic, is a β-coronavirus belonging to the Coronaviridae family. SARS-CoV-2, emerged in 2019, is the third causative agent of severe acute respiratory syndrome also named COVID-19 (CoronaVirus Disease 2019). The two other viruses are SARS-CoV-1 and MERS-CoV emerged in 2003 and 2012, respectively. SARS-CoV-2 is an enveloped virus with a single strand positive RNA genome of about 30 kbases and shares 79% of nucleotide identity with the genome of SARS-CoV (1). Its envelope contains three proteins. i) The “Spike” protein (S) a glycoprotein of 180–200 kDa (2), present as trimers at the surface of the viral particle. It plays a crucial role in the virus entry into target cells following its interaction with ACE2 receptor to induce viral and cell targets membranes fusion (3). ii) The Membrane protein (M), a protein of 25–30 kDa, involved in viral assembly, is the major protein of the envelope. iii) The Envelope Protein (E), is a 8.4-12 kDa polypeptide of 76 to 109 amino-acids (4, 5). It is a small integral viral membrane protein. Inside infected cells, E protein is localized in the RE, Golgi and ERGIC (ER/Golgi intermediate compartments) where it seems to play an important role in the virus assembly and budding (6, 7). In agreement with its role in viral assembly and budding at the RE/Golgi, compartment where are produced the complete viral particles at the end of coronavirus life cycle, its mutation or deletion leads to a substantial decrease in the capacity of viral replication (8, 9). Thus, the important role of E protein, makes it as potential target for antiviral drug molecules and vaccine candidates development (10). Protein-protein interactions were well characterized between E and M proteins, as shown by the presence of E-M complexes at the level of ERGIC in infected cells (11, 12). It is also interesting to note that the expression of these two proteins is sufficient for the formation of VLP (viral like particles) (6, 13). By interfering with protein transport via secretory pathways and with the normal function of the immune system, E protein could act as a pathogenic factor in the immunopathogenesis associated with SARS-CoV, MERS-CoV and SARS-CoV2 (8, 14). In fact, in SARS-CoV infected cells, E protein anchored to a lipid bilayer is able to adopt a structure forming membrane-integral pores, also named viroporin, with a selective activity for cations including, H+, K+, Na+ and Ca2+ (15). At least, the selective permeability to Ca2+ has been reported to be associated with the inflammatory response often observed in ARDS (acute respiratory distress syndrome) (15).

Infection with SARS-CoV-2 is accompanied by deregulation of the control mechanisms of the innate immune response (16, 17). This deregulation is characterized by a delay in the IFN-I and III production and also by an exacerbation of the inflammatory response including, IL-6, TNF-α, IL-1β, IFN-γ but also certain chemokines including CXCL8 (18). In patients developing a critical COVID-19, this dysregulation leads to the establishment of a cytokine storm, a deleterious proapoptotic state for various tissues and organs including the lungs (19–21). SARS-CoV-2 infection also impact the adaptative immune response by affecting the normal physiological functions of antigenic presenting cells (22), but also CD4+, and in a higher degree CD8+ T-cells(23, 24).

Understanding the molecular mechanisms responsible, on one hand, for the control or escape of SARS-CoV-2 detection by innate immune sensors and, on the other hand, for SARS-CoV-2-induced pathological hyper-inflammation are essential steps for the development of effective therapeutic strategies against COVID-19. To achieve this goal, it is important to determine the nature of viral PAMPS and cellular PRRs that are engaged in the course of SARS-CoV-2 infection. According to their biochemical and structural characteristics, PRRs are classified into six different families including: (*i*) Toll-Like Receptors (TLRs); (*ii*) Lectin type C receptors (CLR); (*iii*) scavenger receptors; and (*iv*) the opsonin receptors. In addition to these transmembrane receptors, found on the surface of the cell or in endosomes, cells also express cytosolic and/or nuclear receptors including: (*v*) receptors for nucleic acids, RLR (RIG-I-Like), which recognize the RNAs and cytosolic DNA sensors called CDS, including cGAS, and AIM2-like receptors (ALRs) including AIM2, and (*vi*) NOD type receptors, NOD-Like (NLRs) (25, 26). To date, at least nine PRRs have been reported in the detection of RNA viruses including, TLR7 and 8 (single-stranded RNA), TLR3 (single-stranded RNA), RIG-I and MDA-5 (single and / or double-stranded RNA, di or tri-phosphorylated in 5′), DAI / ZBP-1 (RNA with a Z conformation)(27, 28), receptors forming NLRP3 and NLRP1(29) inflammasomes, as well as helicases of the DDX family including DDX3 which recognizes the RNA of HIV-1(30). More recently, it was advanced that SARS-CoV-2 E envelope protein can be sensed by TLR2 (31), and its expression, as that of its cofactor MyD88, and the induced inflammatory responses seem to increase more importantly in patients with critical severe COVID-19 (32). More interestingly it was shown, in ACE2 transgenic mouse model, that the blockade of TLR2 pathway allowed protection against the disease development and lethality induced by SARS-CoV infection (32).

Considering the important role of innate immune sensors, including TLR2, as potential therapeutic targets in order to alleviate the development of hyper-inflammation and cytokine storm associated with severe COVID-19, in the present study we analysed at molecular level, the interaction between SARS-CoV-2 envelope protein E and human TLR2. Our findings demonstrated that the SARS-CoV-2 E protein interacts with TLR2 receptor in a specific and dose dependent manner in a solid-phase binding assay but also on the cell membrane of TLR2 positive cells, including primary human monocytes and macrophages. Moreover, using HEK-based TLR2 reporter cell lines, we also showed that E protein activates TLR2 signaling pathway that culminate in the activation of NF-κB transcription factor and production of inflammatory cytokines/chemokine including CXCL8.

The finding suggest considering the envelope protein E as a novel target for COVID-19 interventions: (*i*) either by exploring the therapeutic effect of anti-E blocking/neutralizing antibodies in symptomatic COVID-19 patients, or (*ii*) as a promising non-Spike SARS-CoV-2 antigen candidate to include in the development of next generation prophylactic vaccines against COVID-19 infection and disease.

## 2. Materials and Methods

### 2.1 Ethics statement

The use of human cells in this study was approved by the Research Ethical Committee of Haute-Garonne, France. Human Peripheral Blood Mononuclear Cells (PBMC) were isolated from buffy coat of healthy human donors. Buffy coats were provided anonymously by the EFS (Etablissement Français du Sang, Toulouse, France). Written informed consent was obtained from the donors under EFS contract N° 21/PVNT/TOU/INSERM01/2011-0059, according to French Decree N° 2007–1220 (articles L1243-4, R1243-61).

### 2.2 Cells

Human embryonic kidney cell lines stably transfected with TLR2 (HEK-TLR2), TLR4 (HEK-TLR4) and HEK-TLR2-blue and control HEK cell line (HEK-null) were purchased from InvivoGen and cultured in DMEM supplemented with 10 % FCS, 1% of P/S and selections antibiotics according to the manufacturer’s instructions (InvivoGen). Vero E6 and A549 cell lines were cultured in DMEM supplemented with 10% FCS and 1% of P/S.

### 2.3 Virus infection

Primary monocytes-derived macrophages (10^6^ cells) were treated with 0.01 to 1 MOI of the mNeonGreen SARS-CoV-2. This recombinant reporter SARS-CoV-2 developed by Pei Yong Shi et al (33) was obtained from World Reference Center for Emerging Viruses and Arboviruses (WRCEVA).

### 2.4 Isolation of human monocytes

PBMCs were isolated from buffy coats of healthy blood donors (from Etablissement Français du Sang [EFS], Toulouse) and monocytes were separated from lymphocytes by positive selection using magnetic cell sorting technique according to the manufacturer’s instructions (Miltenyi Biotec) and as described (34).

### 2.5 Generation of monocyte-derived macrophages

To allow differentiation of monocytes into monocyte-derived macrophages, monocytes were cultured in DMEM medium (Invitrogen) supplemented with 10% fetal calf serum (FCS) 100 IU/ml penicillin, 100 μg/ml streptomycin, 10 ng/ml GM-CSF, and 10 ng/ml MCSF. After 3 days of culture, cells were stimulated by the same amount of GM-CSF and M-CSF and cultured for additional 4 days before their use in our experiments as differentiated macrophages.

### 2.6 Chemical products, Proteins, and Antibodies

PAM_2_CSK_4_, PAM_3_CSK_4_, LPS-RS were purchased from InvivoGen. Recombinant soluble E protein from SARS-CoV-2 was purchased from Clinisciences. GST, GST-Nef and the corresponding antibodies were produced in our laboratory. Soluble recombinant TLR2 was purchased from R&D systems. Anti-TLR2 and anti-TLR4 monoclonal antibodies were obtained from eBioscience. Anti-Phospho-P65 and anti-total P65 were purchased from cell Signalling. Bay11-7082, SB202190, PD98059 and RO318220 were purchased from Calbiochem.

### 2.7 Interaction of E protein with TLR2 in a solid phase assay

The binding of the recombinant E-GST protein with TLR2 was tested in a solid phase assay. Briefly, 100 μL of recombinant soluble TLR2 (R&D systems) at 1 μg / ml are coated during 2 hours at room temperature in 96-well plates. After 1 hour of saturation with 300 μL of PBS containing 5% non-fat milk and 5 washes with PBS-Tween 0.05%, 100 μL of different concentrations of the soluble protein E-GST (1ng-1000 ng/ml) are added to each well. After 1 hour of incubation at 37 °C, 5 washes were performed with PBS-0.05% Tween. Then, the detection of TLR2-E-GST complexes were performed by an additional incubation during 1 hour at room temperature with 100μl of a rabbit anti-GST sera previously diluted at 1/500 in PBS-tween 0.05% containing 5% non fat milk. After 5 further washes, the complexes TLR2-E-GST-anti-GST were labeled by 1 hour incubation at room temperature with 100 μl of anti-rabbit IgG antibodies coupled to horseradish peroxidase in PBS-tween 0.05% containing 5% non-fat milk (DAKOTA). After a last 5 washes with PBS-tween 0.05%, TLR2-E-GST-anti-GST-anti-rabbit-IgG-peroxydase complexes were revealed by the addition of 100 μL of TMB substrate (Tetramethylbenzidine). After 15 to 30 min incubation, the peroxidase reaction was stopped with 50 μL of sulfuric acid (2N) and then the optical density was read at 450/570 nm.

### 2.8 Inhibition assay of E-TLR2 interaction

The specificity of E-TLR2 interaction was evaluated in a solid phase binding assay as described above except that various amounts of PAM_2_CSK_4_, PAM_3_CSK_4_ were added to rTLR2-precoated wells during 1 hour before adding a constant amount of soluble E protein (200ng).

### 2.9 Flow cytometry analysis

Monocytes (10^6^) were incubated with GST or GST-E SARS-CoV-2 protein at 0.1-10μg/ml for 1 hour at 37 °C in PBS, BSA 0.5%, NaN_3_ 0.05%. Then, cells were washed 3 times with PBS, BSA 0.5%, NaN_3_ 0.05% to remove unbound proteins. Cells were stained with and anti-GST-Alexa 488 (Catalog # A-11131, ThermoFisher, 1/2000) during 1 hour at room temperature and washed 3 times with PBS, BSA 0.5%, NaN_3_ 0.05%. Then cells were fixed with PFA 4%. Data were acquired using FACSCalibur (BD).

### 2.10 Microscopy analysis

The analysis of the binding of E protein to macrophages was analyzed by microscopy. To this end macrophages (10^6^) were incubated with GST or GST-E SARS-CoV-2 protein at 10μg/ml for 1 hour at 37°C PBS, BSA 0.5%, NaN_3_ 0.05%. Then, cells were washed 3 times with PBS, BSA 0.5%, NaN_3_ 0.05% to remove unbound proteins. Then cells were washed 3 times with PBS, stained with Hoechst, and anti-GST-Alexa 488 (1/500) during 1 hour at room temperature and washed 3 times with PBS, BSA 0.5%, NaN_3_ 0.05%. Finally, cells were fixed with PFA 4% before imaging. Images were acquired using EVOS M700 (Invitrogen) at 40x magnification.

### 2.11 Cell based biological assays

Primary human monocytes or macrophages cells (10^6^ cells) or HEK-null, HEK-TLR2 or HEK-TLR4 cell lines (2,5. 10^5^ cells) were plated in 24 well plates and treated by E protein or PAM_3_CSK_4_ and PAM_2_CSK_2_ as positive controls at the indicated concentrations. Untreated cells were used as negative controls. To block TLR2, anti-TLR2 were added in cell culture medium 1 hr before treatment with E protein. To inhibit cell signaling pathways, cells were incubated with chemical inhibitors 30 minutes before treatment with E protein. To inhibit the binding of E protein to cell membrane TLR2, E protein (at 200ng/ml) was preincubated with rTLR2 (20 ng/ml) during 1 hour at RT, before being added to HEK-TLR2 cells. Cell supernatants were collected 18hrs after E-treatment and frozen at −20°C before further analysis.

### 2.12 Phosphorylation analysis of NF-kB P65 subunit and Western blot analysis

HEK-TLR2 cells (2,5.10^5^ cells) were treated during 30 or 60 min with E protein (1 μg/ml) or, with GST (1μg/ml) or PAM_3_CSK_4_ (10 ng/ml) as negative and positive controls respectively. Then, cells were lysed and prepared for immunoblot as previously described (35).

### 2.13 NF-kB assay using HEK-TLR2-Blue

The capacity of E protein to activate NF-kB was tested by using HEK-TLR2-blue (InvivoGen). In this assay, HEK-TLR2 cells stably transfected with SEAP (secreted embryonic alkaline phosphatase) gene under the control of NF-κB promoter were plated at 2,5.10^5^ cells per well in 24 well plates one day before the experiment. The following day cells were treated by E protein in cell culture medium at the indicated concentration. 18 hrs after treatment, supernatants were collected and quantification of SEAP was performed according to manufacturer’s instructions (InvivoGen).

### 2.14 CXCL8 quantification by ELISA

Cells were stimulated with various amount of E protein (1-1000ng/ml). After 18 hours of stimulation at 37°C, supernatants were harvested and stocked at −20°C until CXCL8 quantification by ELISA kits according the instructions of the manufacturer (R&D system).

### 2.15 Statistical analyses

Statistical analysis was performed using GraphPad Prism software v.5. All results are expressed as means +/− SD. All experiments were performed a minimum of three times. Differences in the means for the different groups were tested using one-way ANOVA followed by Bonferroni post hoc test. A p-value <0.05 was considered statistically significant. Statistical significance comparing different groups is denoted with * for p < 0.05, **p < 0.01, ***p < 0.001, ns non-significant.

## 3. Results

### 3.1 SARS-CoV-2 E-envelope protein interacts directly and physically with TLR2

In order to analyse the capacity of SARS-CoV-2 envelope protein E to interact with TLR2 at a molecular level, we tested in a solid phase assay the binding of various amounts of E protein (1 ng/ml-1000 ng/ml) to a constant amount of pre-coated human recombinant TLR2 (1 μg/ml). The obtained results depicted in **Figure 1** showed that E protein binds in a dose-dependent manner to TLR2. In contrast, no significant binding to TLR2 was observed when the experiment was performed with GST, instead of E protein (**Figure 1A**). Further, we also showed that E protein, but not GST used as control, is also able to bind to human primary monocytes when analysed by flow cytometry (**Figure 1B, C**) and to human primary macrophages when analysed by microscopy (**Figure 1D**).

**Figure 1:**
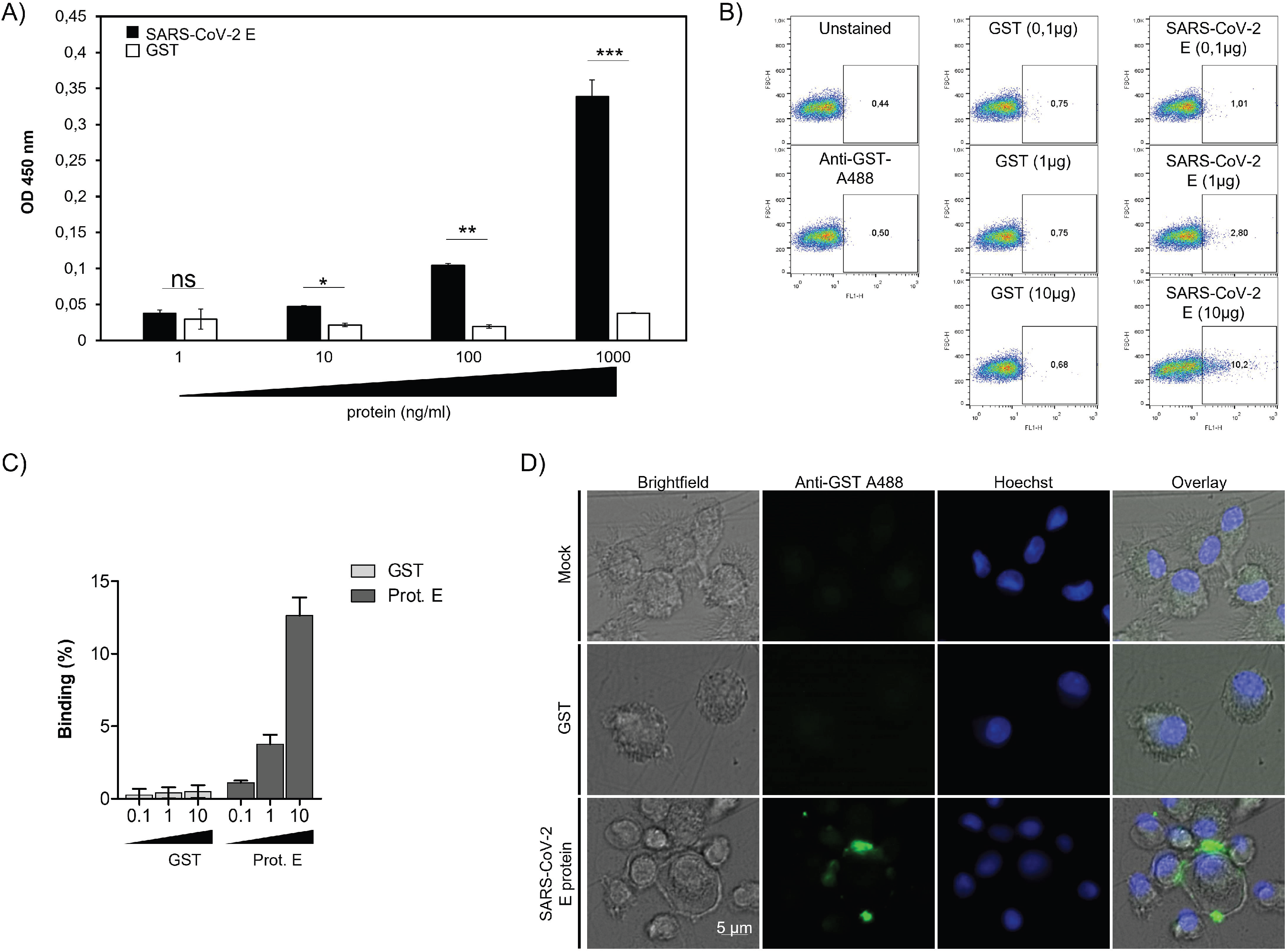
Binding of SARS-CoV-2 protein to human TLR2: (**A**) Soluble recombinant human TLR2 (100 μl at 1 μg/ml) were coated in 96 plates. After saturation, various amounts of E-GST protein (1 ng/ml-1000 ng/ml) were added for 2 hours at 37°C. TLR2 E-GST complexes were revealed by a solution of anti-GST-sera follow by anti-anti-GST conjugated to HRP. (**B**) Primary human monocytes were incubated with 0,1 to 10 μg/ml of GST or GST-E SARS-CoV-2 protein. Cells were stained with anti-GST (1/1000). Data were acquired using FACScalibur. One representative experiment is shown. (**C**) Quantification of SARS-CoV-2 E protein or GST control binding to human monocytes out of from 3 different experiments acquired on FACScalibur. (**D**) Primary human macrophages were incubated with 10 μg/ml of GST or GST-E SARS-CoV-2 protein. Cells were stained with anti-GST (1/500). Images were acquired using EVOS M700 microscope.

Altogether, these results demonstrate that the SARS-CoV-2 envelope protein is able to interact physically, in a dose-dependent manner with the human soluble recombinant TLR2 but also with cell membrane TLR2 expressed at the surface of primary human monocytes and macrophages.

### 3.2 PAM_2_CSK_4_ and PAM_3_CSK_4_ antagonise SARS-CoV-2 E protein binding to TLR2

PAM_2_CSK_4_, a synthetic diacylated lipopeptide ligand of TLR2/TLR6, and PAM_3_CSK_4_, a synthetic triacylated lipopeptide ligand of TLR2/TLR1 have been historically characterized as the first identified ligands of TLR2. Thus, in order to characterise the specificity of the interaction of E protein with TLR2, we evaluated the capacity of these two ligands PAM_2_CSK_4_ and PAM_3_CSK_4_ to inhibit E-TLR2 interaction. To this end, the experiment was performed as described in figure 1, by using a constant concentration of E protein (200ng) but in presence of escalating amounts of PAM_2_CSK_4_ or PAM_3_CSK_4_ (0.1μM to 10μM). Both ligands inhibited E-TLR2 interaction in a dose dependent-manner (**Figure 2A-B**). However, only a partial inhibition, exceeding 50%, was obtained with PAM_2_CSK_4_ and PAM_3_CSK_4_ used at 10μM. Thus, these characterizations demonstrate that E protein-TLR2 interaction is specific as demonstrated by the capacity of PAM_2_CSK_4_ and PAM_3_CSK_4_ to inhibit this interaction.

**Figure 2:**
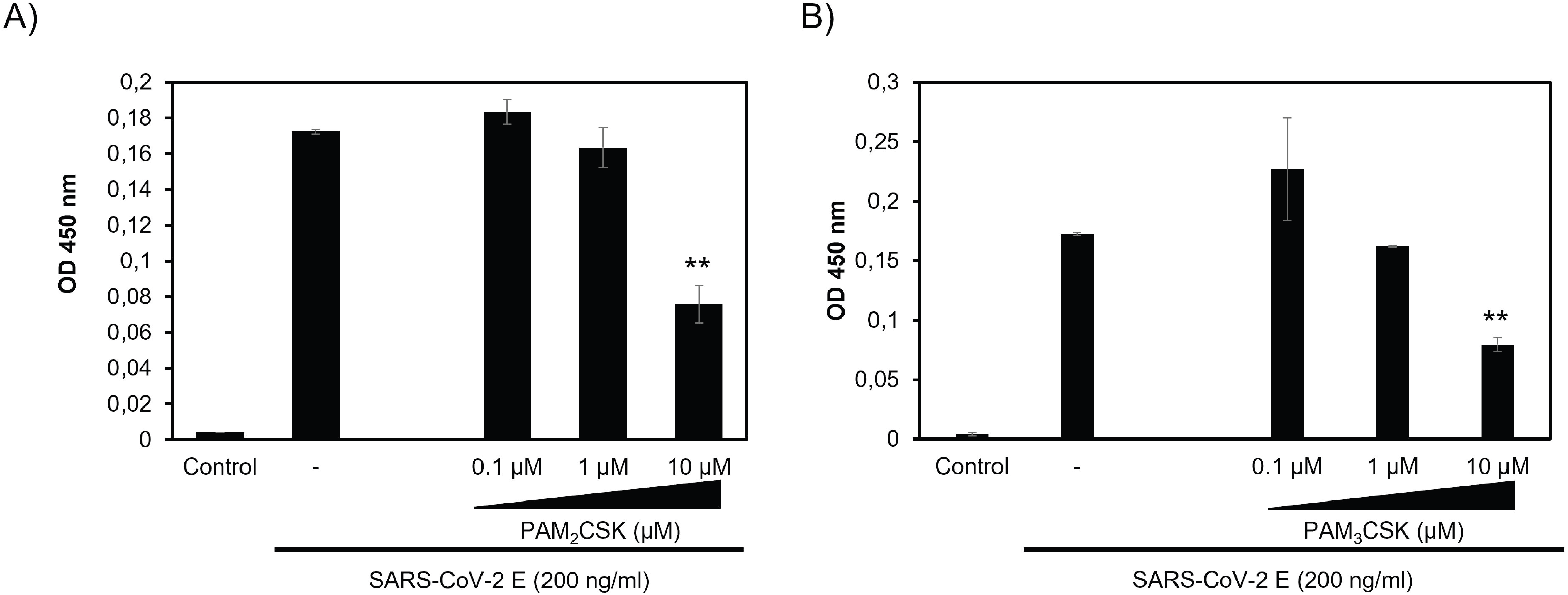
PAM_2_CSK_4_ and PAM_3_CSK_4_ interfere with SARS-CoV-2 E binding to TLR2: The specificity of E-TLR2 interaction was evaluated by testing the capacity of TLR2 ligands (**A**) PAM_2_CSK_4_ (0.1-10 μM) and (**B**) PAM_3_CSK_4_ (0.1-10 μM) to inhibit this interaction.

### 3.3 SARS-CoV-2 E Protein stimulates the production of CXCL8 inflammatory chemokine by recruiting TLR2 pathway

In order to study the biological consequences of E-TLR2 interaction, we tested the capacity of E protein to stimulate the production of CXCL8 in a HEK cell lines-based assay using cells stably transfected with the human TLR2 (HEK-TLR2) or TLR4 (HEK-TLR4) receptors or HEK-null (transfected with empty plasmid). As previously shown by our group, activation of TLR-dependent pathway in HEK cells lines stimulates the production of measurable amount of CXCL8 chemokine, while other TLR-dependent cytokines including TNF-α, IL-6, IL-10 where barely detectable. As consequence, the production of CXCL8 by HEK cells lines was used as a marker of TLR response (36). Results presented in figure 3A show that E protein from SARS-CoV-2 (200ng/ml) stimulates the production of CXCL8 in TLR2-expressing HEK cell lines, while GST or GST-Nef, two unrelated SARS-CoV-2 gene products, used as controls, do not stimulate significant production of CXCL8 when used at concentrations up to 1 μg/ml (**Figure 3A**). As additional controls, no production of CXCL8 was obtained in the supernatants of unstimulated HEK-TLR2 cell line, while a clear production of CXCL8 was produced following the stimulation by the synthetic ligand of TLR2 PAM_3_CSK_4_ (**Figure3 A**). The specificity of the activation of TLR2 pathway by E protein was further demonstrated by showing that E protein induced the production of CXCL8 in a dose-dependent manner, with the lowest amount of E protein giving a detectable CXCL8 production being around 10 ng/ml (**Figure 3B**). The specificity of E-TLR2 pathway activation was further supported by the fact that no CXCL8 production was obtained in HEK-null cell (**Figure 3C**) nor in HEK-TLR4 (**Figure 3D**) cell lines. This latter control also demonstrated the absence of endotoxins contaminants in our recombinant E protein as demonstrated by the absence of any activation of TLR4 pathway (**Figure 3C-D**).

**Figure 3:**
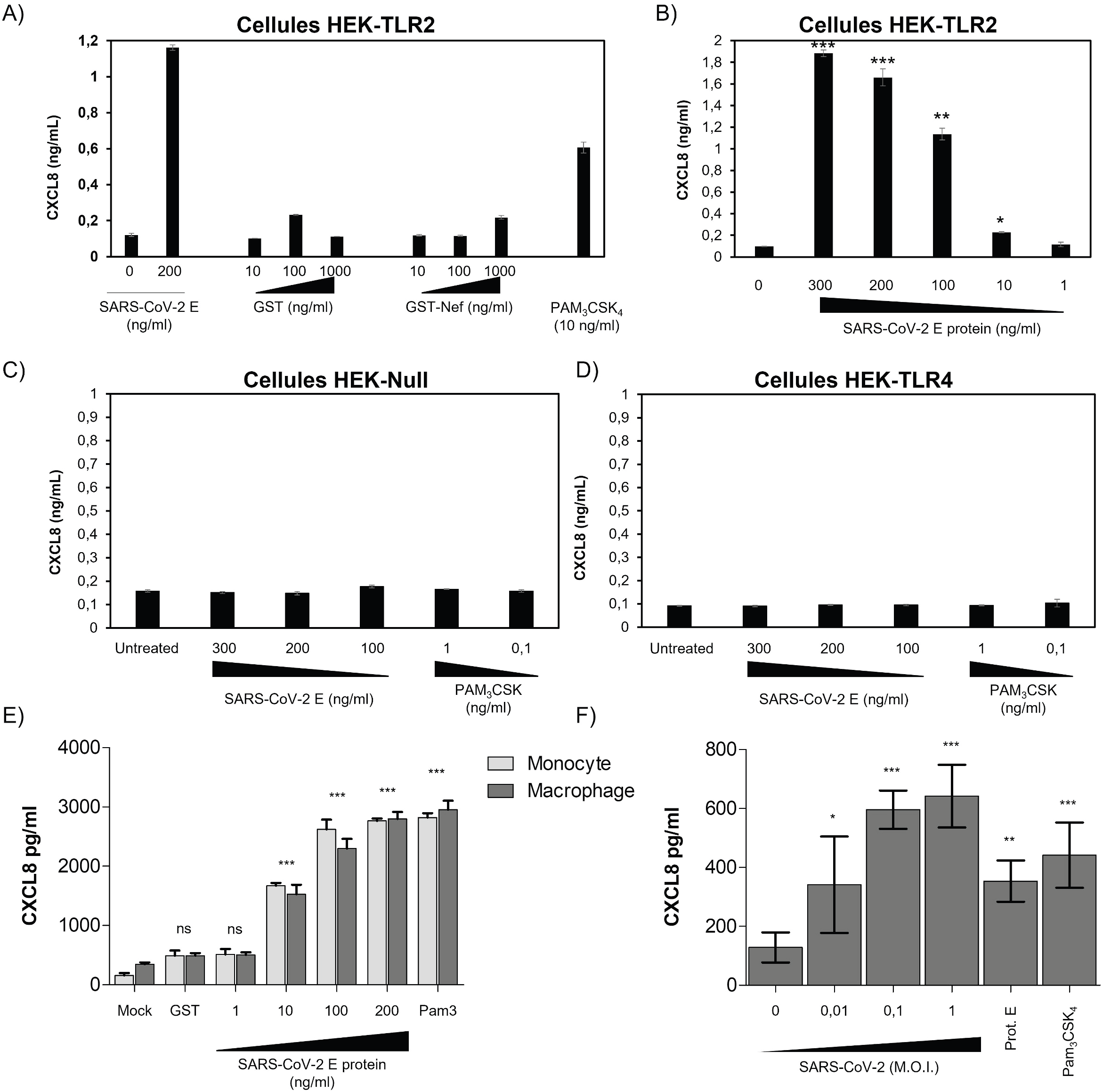
E protein stimulate the production of CXCL8 chemokine in a TLR2-dependent manner: (**A**) HEK-TLR2 cell line were stimulated with E protein (200ng/ml), GST (10-1000ng/ml) or GST-nef (10-1000ng/ml), or PAM_3_CSK_4_ (10 ng/ml). CXCL8 chemokine production in the cell supernatants was quantified by ELISA. (**B**) Production of CXCL8 in cell supernatants of HEK-TLR2 cells stimulated by escalating concentrations of E protein (1-300ng/ml). (**C-D**) Production of CXCL8 in cell supernatants of HEK-null (**C**), HEK-TLR4 (**D**), cell lines stimulated with E protein (1-100ng/ml) or with PAM_3_CSK_4_. (**E**) Primary human monocytes and macrophages were stimulated with E protein (1-200ng/ml). Stimulation with GST (200 ng/ml) or PAM_3_CSK_4_ (1000 ng/ml) were used as negative and positive control respectively. After 20h of treatment cell supernatant was collected and CXCL8 chemokine production in the cell supernatants was quantified by ELISA. (**F**) Primary human macrophages were infected with NeonGreen SARS-CoV-2 virus (MOI 0.01-1). Stimulation with SARS-CoV-2 E protein (10 ng/ml) or PAM_3_CSK_4_ (10 ng/ml) were used as positive control. After 20h of treatment cell supernatant was collected and CXCL8 chemokine production in the cell supernatants was quantified by ELISA.

Taking into account the data obtained with HEK-TLR2 cell line, we tested the capacity of E protein to activate the production of CXCL8 in human primary monocytes and macrophages. To this end, human monocytes and macrophages were stimulated by increasing concentrations of E protein (1 ng/ml to 200 ng/ml) for 20 hours and CXCL8 was quantified by ELISA as described above. The obtained results showed that E protein stimulated the production of CXCL8 in both primary human monocytes and macrophages (**Figure 3E**).

In line with these results, we also showed that treatment of primary human macrophages, during 20 hours, with infectious SARS-CoV-2 viral particles also resulted in the production of CXCL8 in cell supernatants (**Figure 3F**). It is interesting to note that macrophages do not show any signs of viral replication, as shown by the absence of NeonGreen fluorescence in macrophages when infected with the recombinant mNeonGreen SARS-CoV-2, used at 0.01, 0,1 and 1 MOI (**Supplementary Figure S1**). As positive control, we showed that the treatment of VeroE6 cells with the recombinant mNeonGreen SARS-CoV-2 resulted in a clear infection of these cells (**Supplementary Figure S1**).

Altogether, our results showed that SARS-CoV-2 E protein, by recruiting, at least, TLR2 pathway stimulated the production of CXCL-8.

### 3.4 The stimulation of CXCL8 production by SARS-CoV-2 E protein is reversed by soluble rTLR2 and anti-TLR2 antibodies

The specificity of the recruitment of TLR2 pathway by E protein was further characterized in complementary assays using either soluble recombinant TLR2 (rTLR2) or anti-TLR2 blocking antibodies. The results show that incubation of rTLR2 with E protein before stimulation of HEK-TLR2 cells inhibit by about 50% the capacity of E protein to stimulate TLR2-response as measured by the production of CXCL8. Importantly, no significant CXCL8 production was obtained with rTLR2 alone (**Figure 4A**). Interestingly, this latter result also indicates the absence of endotoxins in the used preparation of rTLR2. In agreement with the effect of rTLR2 we showed that anti-TLR2 antibodies used at 5μg/ml, inhibits by about 60% E-induced CXCL8 production (**Figure 4B**) while only a moderate inhibition, less than 30%, was observed by the use of anti-TLR4 antibodies (5μg/ml), used as isotype controls. We also tested the effect of LPS-RS a specific antagonist of TLR4. Used at 10 μg/mL, LPS-RS induced a modest inhibition of E2 induced CXCL8 production of about 18% (**Figure 4C**). These moderate inhibitions may be caused by the steric hindrances caused by the presence of anti-TLR4 antibodies and LPS-RS.

**Figure 4:**
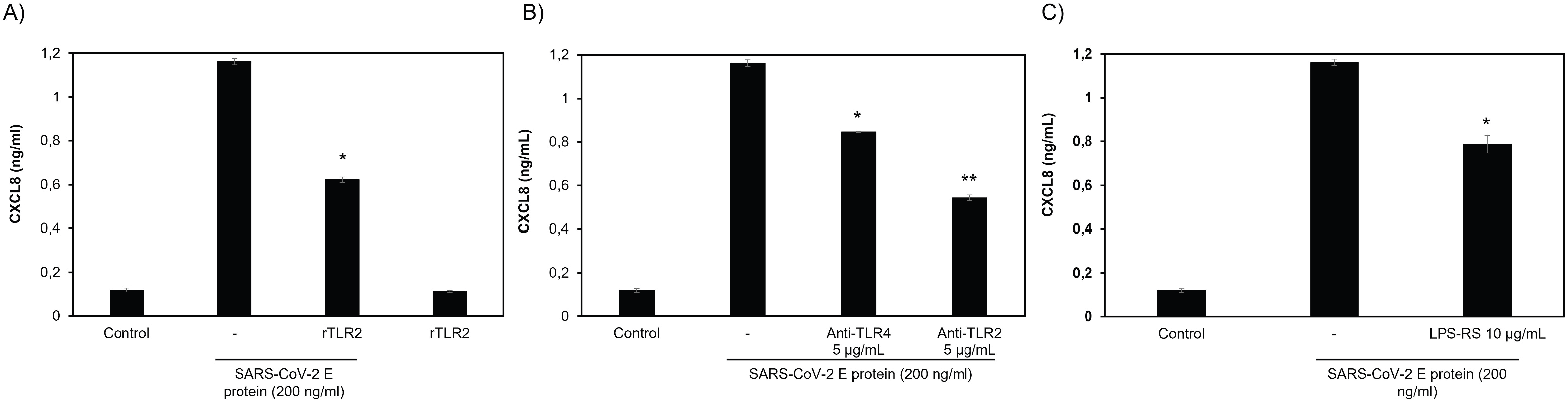
Inhibition of E-induced CXCL8 production by soluble recombinant TLR2 and anti-TLR2 antibodies: HEK-TLR2 cell line were stimulated with E protein (200ng/ml) in the presence or absence of recombinant TLR2 (20 ng/ml) (**A**), anti-TLR2 or anti-TLR4 antibodies (**B**) or LPS-RS (10 μg/ml) as control (**C**). HEK-TLR2 cells were also treated with recombinant TLR2 (20 ng/ml) alone as control (**A**). After 20h of treatment cell supernatant was collected and CXCL8 chemokine production in the cell supernatants was quantified by ELISA.

Altogether, these results confirm the recruitment of TLR2 pathway by E protein as demonstrated by the capacity of soluble recombinant TLR2 and anti-TLR2 antibodies to strongly block the production of CXCL8 chemokine production in HEK-TLR2 cells stimulated by SARS-CoV-2 E protein.

### 3.5 SARS-CoV-2 E protein activates NF-kB as a signature of the recruitment of TLR2 pathway

Activation of all TLR pathways leads to activation of the NF-kB. NF-kB is an important transcription factor greatly implicated in the control of the expression of cytokines genes that are involved in the immune and inflammatory responses (37, 38). The analysis of the *CXCL8* promotor element sequence highlights the presence of NF-kB binding site. NF-kB, a REL family member is composed of hetero or homodimers of 5 subunits including RelA/p65, c-Rel, RelB, p50 and p52 (39). At the inactivated state heterodimeric NF-kB is present in the cytoplasm in association with its inhibitor IkB. In order to be activated, NF-kB must be phosphorylated on its subunits p65 and p50, but also on its inhibitor subunit IkB, thus leading on one hand, to the nuclear translocation of P65 into the nucleus, where it binds on NF-kB sites at the CXCL8 promotor element sequence, and on the other hand, on the dissociation, ubiquitinilation and proteasomal degradation of IkB (38). Here, in our study, the effect of E protein on the activation of NF-kB was evaluated by monitoring its effect on the phosphorylation of the p65 subunit. To this end, HEK-TLR2 cells were stimulated during 30 or 60 min with E protein (1 μg/ml) or with GST or PAM_3_CSK_4_ as negative and positive controls respectively. Both at 30 and 60 min post stimulation, E protein leads to the phosphorylation of p65 (**Figure 5A lanes 3 and 4**). Only a small phosphorylation of p65 was observed in unstimulated cells (**Figure 5A lane 2**) and in cells stimulated with GST protein (**Figure 5, lanes 5 and 6**). As expected, a strong phosphorylation was obtained following stimulation with PAM_3_CSK_4_ (**Figure 5A lane 7**).

**Figure 5:**
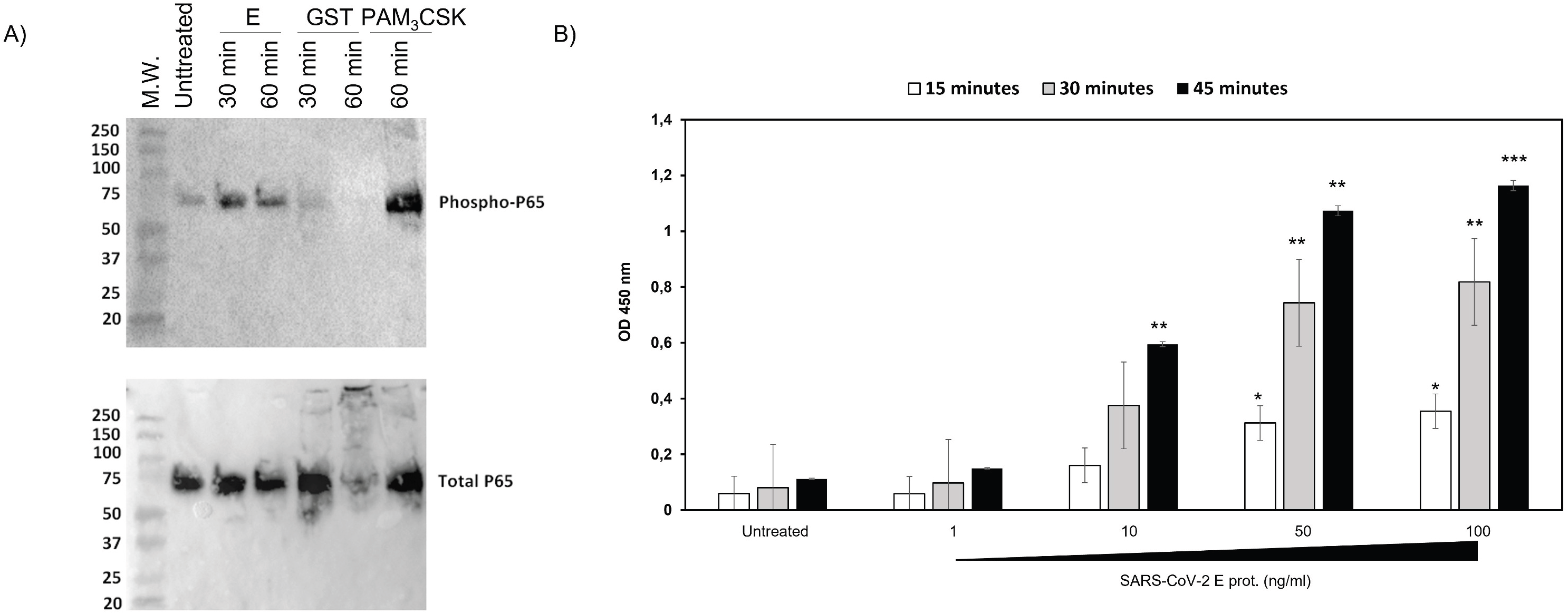
E protein stimulates the activation of NF-kB: (**A**) HEK-TLR2 cells were stimulated with E protein, GST or PAM_3_CSK_4_ during 30 or 60 min. Phosphorylation of P-65 was analysed by SDS-PAGE and western blot by using specific anti-phospho-P65 (upper panel) or anti-total p65 antibodies (lower panel). (**B**) HEK-TLR2 cell line, stably transfected with SEAP (secreted embryonic alkaline phosphatase), were treated with escalating concentrations of E protein or with PAM_3_CSK_4_ during 15, 30 and 45 min and SEAP activity was quantified in the cell supernatants.

Then, the effect of E protein on the activation of NF-kB was further characterized in a more functional assay, based on the evaluation of the capacity of E protein to transactivate the expression of the gene product of SEAP soluble protein placed under the control of NF-kB inducible promotor. To this end, HEK-TLR2 cell line stably transfected with SEAP gene under the control of NF-kB, were stimulated with various amount of E protein (1ng-100 ng/ml) during 15 min, 30 min or 45 min. The expression of the enzymatic activity of soluble secreted SEAP protein was then measured in the cell supernatants. The obtained results depicted in **Figure 5B** clearly showed a positive presence of SEAP enzymatic activity since 15 min of stimulation with 10ng/ml of E protein. This enzymatic activity increased in time and in a dose-dependent manner at 30 min and 45 min following stimulation with the highest doses of 50 ng/ml and 100 ng/ml (**Figure 5 B**). As negative control, no significant SEAP enzymatic activity was observed in supernatants of unstimulated HEK-TLR2 cells (**Figure 5 B**).

Altogether, these results showed that SARS-CoV-2 E envelope glycoprotein is able to recruit and engage TLR2 pathway leading to the activation of the transcription factor NF-kB as demonstrated by the phosphorylation of p65 NF-kB subunit and the transactivation of SEAP gene under the control of NF-kB promotor site.

### 3.6 SARS-CoV-2 E protein activation of CXCL8 production is dependent on NF-kB pathway

Then we wanted to evaluate the role of NF-kB in the control of CXCL8 production in response to E stimulation in HEK-TLR2. To this end, HEK-TLR2 cell line cells were previously treated during 60 min with various non-toxic concentrations (1–10 μM) of NF-kB inhibitor Bay11-7082 before stimulation with E protein (200 ng/ml). After 18 h of culture, CXCL8 production was quantified in cell supernatants. A dose-dependent inhibition of CXCL8 production was obtained in the presence of Bay11-7082 demonstrating the crucial role of the transcription factor NF-kB in the control of gene expression of CXCL8 chemokine (**Figure 6A**).

**Figure 6:**
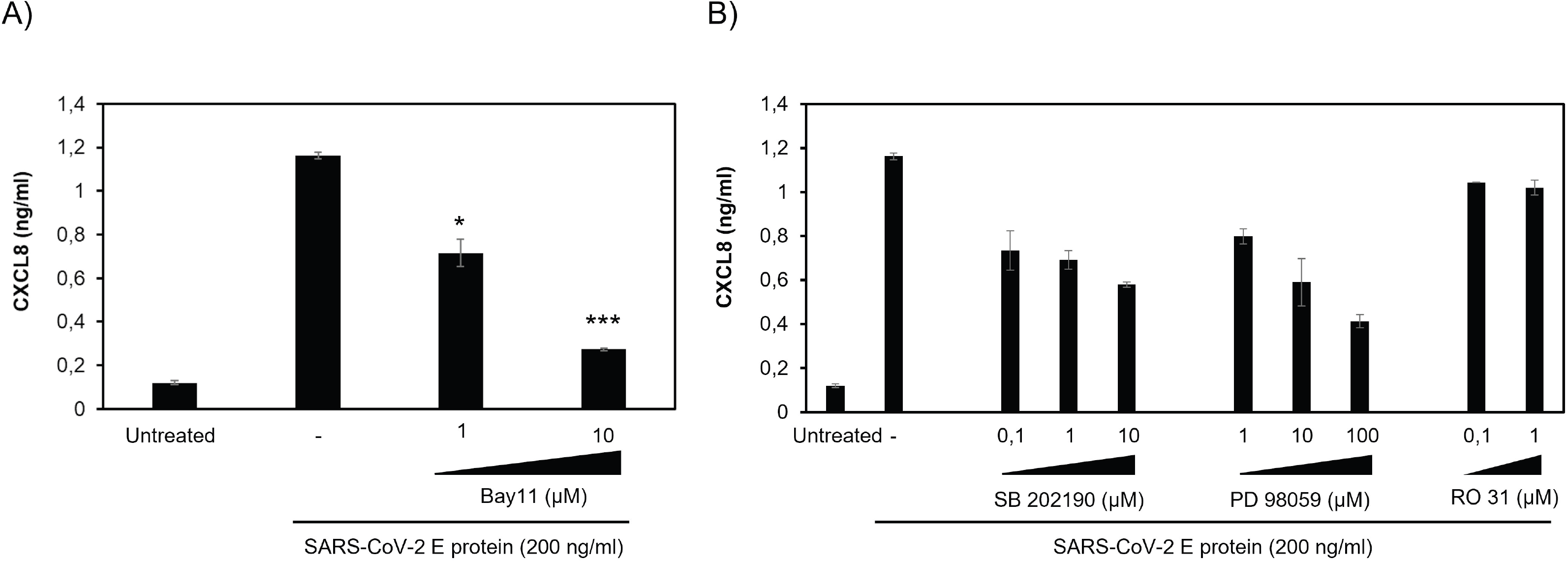
inhibition of E-induced CXCL8 production by NF-kB inhibitor but not by P38 and ERK1/2 MAPkinases and PKC inhibitors: (**A**) inhibition of E induced-CXCL8 chemokine in the presence of NF-kB inhibitor. HEK-TLR2 cells were stimulated with E protein (200ng/ml) in the presence of the chemical inhibitor of NF-kB Bay11 used at 1 and 10 μM. Production of CXCL8 in cell supernatants was quantified by ELISA. (**B**) HEK-TLR2 cells were previously treated with P38 MAP kinase inhibitor SB202190 (0.1-10μM), ER11/2 inhibitor MAP kinase PD 98059 (1-100μM) or PKC inhibitor RO318220 (0.1-10μM) during 1 hour, before stimulation with E protein (200ng/ml). Produced CXCL8 in cell supernatants was quantified by ELISA.

In addition to NF-kB, the promotor element sequence of CXCL8 gene contains also binding sites for additional transcription factors, including AP-1 (activating protein), CREB (cAMP response element binding protein), C/EBP (CAAT/enhancer-binding protein), CHOP (C/EBP homologous protein) (40) and C/EBP beta (also named NF-IL-6) (41). While NF-kB is crucial for the gene expression of CXCL8, the other transcription factors, as AP1 and CREB seem to play a secondary role by acting on the stability of mRNA and synergy action with NF-kB on the expression of CXCL8 gene, thus contributing to allow an efficient production of CXCL8 gene product (41, 42). The MAPkinases, including P38MAPK and ERK1/2 MAPK has been reported to participate in the activation of AP1 and CREB and thus, indirectly via AP1 and CREB, in the contribution of the increased expression of CXCL8 gene product. Taking into account these contributions, we tested the effect of the inhibition of P38 MAP and ERK1/2 on the production of CXCL8 following activation of HEK-TLR2 cell line by E protein. To this end, HEK-TLR2 cells were previously treated during 60 min by a non-toxic concentrations of SB202190 (0.1-10 μM) and PD98059 (1-100 μM) as inhibitors of P38MAPK and MAPK ERK1/2 respectively before treatment with E protein at 200ng/ml. Both inhibitors exhibit a partial inhibitory effect reaching respectively, 55% and 70 % of inhibition by SB202190 and PD98059 when used at the highest concentrations (**Figure 6B**).

Because P38 and ERK1/2 are activated downstream of PKC, a large family of serine /threonine kinase, we also evaluated the effect of PKC on the production of CXCL8 by E stimulated HEK-TLR2 cells. To this end, cells were previously treated with various concentrations of R0318220, an inhibitor of all PKC isoforms, before stimulation with E protein at 200 ng/ml and quantification of CXCL8 in cell supernatants as described above. No evident inhibition was obtained in the presence of the PKC inhibitor Ro318220 used at 0.1 and 1μM (**figure 6B**). However, the apparent inhibition observed at 10 μM of the inhibitor is further related to the cytotoxic effect of 10μM concentration of RO318220 as evaluated by the cytotoxicity assay measuring LDH release, a signature of cell death (data not shown).

Taken together, our results demonstrated the direct physical interaction between the E envelope protein of SARS-CoV-2 and the TLR2. This interaction engages the activation of TLR2 pathway leading to the activation of the transcription factor NF-kB which seems to play, in contrast to ERK1/2 and P38 MAP kinases, a major role in the production of CXCL8 chemokine.

## 4. Discussion

Recent work by Zheng and colleagues provided genetic evidence that TLR2 pathway contributes to overwhelming production of inflammatory cytokine production (particularly TNF-α, IL-6, IFN-γ) during infection by SARS-CoV-2 and other β-coronaviruses, following recognition of E envelop protein (31). In light of this recent finding, our study provides further characterization of E – TLR2 interaction. Specifically, our results demonstrate that SARS-CoV-2 E envelope protein interacts physically in a dose-dependent manner with soluble recombinant TLR2 receptor but also with cell membrane TLR2 of primary human monocytes and macrophages. Additionally, our findings show that E protein from SARS-CoV-2 activates TLR2 pathway leading to the activation of the transcription factor NF-kB which seems to play a major role in the production of CXCL8 chemokine, in contrast to ERK1/2 and P38 MAP-kinases whose inhibition only results in partial inhibition of CXCL8.

TLR2 was originally described to recognize ligands from bacterial origins (43–46) that include diacyl and triacylglycerol moieties, proteins and polysaccharides. However, it is currently assumed that recognition of TLR2 is not limited to bacterial ligand but concern a broader set of molecules including viral proteins (review in (47)). These TLR2 viral ligands include Glycoprotein B of Cytomegalovirus, hepatitis C core and NS3 Protein, and hemagglutinin (H) of measles virus (47). Thus, E protein from SARS-CoV-2 extend the list of viral TLR2 ligands. The diversity of molecules recognized by the receptor TLR2 may be licensed by its capacity to form heterodimer with TLR1, TLR6 or TLR10 and to benefit from the help of additional cofactors including CD14 and CD36 (47). However, the involvement of TLR2 in the interaction with E protein raises a number of questions. Crystallographic studies of the complex between TLR2/TLR1 and its tri-acylated lipopetides ligands PAM_3_CSK have allowed to determine the sites of interaction between TLR2/TLR1 with their ligands PAM_3_CSK (48). The structures of the lipopeptide TLR2 ligands, PAM_3_CSK and PAM_2_CSK contain 3 and 2 lipid chains respectively. By interacting with the hydrophobic pocket of TLR2, these lipid chains allow heterodimerization of TLR2/TLR1 by PAM_3_CSK and TLR2/TLR6 by PAM_2_CSK and the recruitment of downstream adapters including Mal/Myd88, thus allowing to the activation of the TLR2 pathway (48). It is therefore important to question how the E protein of SARS-CoV-2, which does not have a lipid tail, can interact and activate the TLR2 pathway. Indeed, the analysis of the primary structure of the E protein reveals two hydrophilic regions in the N and C terminal parts of the molecule separated by a large hydrophobic domain which could present an affinity for the hydrophobic pocket of TLR2. In addition, it has been reported that E protein also exists in the form of homo-oligomeric multimers (15, 49, 50) which by interacting with the hydrophobic pockets of TLR2, TLR1 and TLR6 could bridge the formation of heterodimers of TLR2/TLR1, or TLR2/TLR6 and even of homodimers of TLR2/TLR2. Thus, further structural research studies are needed to confirm these hypotheses. Our data showing that PAM_3_CSK and PAM_2_CSK synthetic ligands interfere with E-TLR2 binding, suggest that E protein and PAM_2_CSK/PAM_3_CSK bind TLR2 on partially overlapping sites.

In our study, we demonstrated a direct physical binding of E protein to TLR2 in a solid phase binding assay. However, this assay is not informative about the functionality of this interaction, nor it does not indicate if it induced structural rearrangements or oligomerizations of TLR2. But, our findings showing that E protein is also able to bind to cell membrane TLR2 of primary monocytes and macrophages, to activate the transcription NF-kB and to stimulate the production of CXCL8 chemokine represent strong arguments in favour of the capability of E protein to recruit and engage TLR2 pathway.

Activation of TLR2 light a diverse number of intracellular signalling pathways that culminate in transcription of several immunity related genes including pro-inflammatory cytokines and chemokines with important role in shaping innate and adaptive immune response, as well as tissue homeostasis. Our data show that E protein activates NF-kB and demonstrates that this activation is essential for CXCL8 production in HEK-TLR2 cell line. The partial inhibition obtained in the presence of P38 and ERK1/2 MAPkinases inhibitors is in line with the secondary role of these pathways, involved in the activation of the AP1 and CREB transcription factors in the CXCL8 gene expression (41, 42).

Activation of TLR pathway by viruses play a mitigated role and triggers either immune protection or pathogenesis of infections (51, 52). For example, the use of animal model elegantly exemplifies that TLR7-dependent type I interferon production by plasmacytoïd dendritic cells (pDCs) confers protection against mouse hepatitis virus (MHV) viral infection (53). Similarly, type I interferon production was also observed following pDCs interaction with SARS-CoV (53) and SARS-CoV-2 (54). Accordingly, in order to escape from TLR-mediated immunity, viruses have developed several strategies to interfere with signal transduction downstream of TLR pathways (52, 55). In contrary, dysregulated activation of TLR pathway has been associated with enhanced pathogenesis. This is the case for TLR4 pathway which is involved in pathogenesis of IAV, EBOV, and DENV infections while treatment with TLR4 antagonists (Eritoran) reduced cytokine/chemokine production and alleviate disease symptoms (56). Other viruses are taking advantage of TLR pathway to their own benefits. Our group and other have shown that HIV-1, through its Tat protein, activates TLR4 pathway leading to the upregulation of several immunosuppressive factors including IL-10, PD-L1 and IDO-1 (35, 57–59). This is also the case for measles virus which subvert TLR2 pathway by its hemagglutinin (H) protein in order to upregulate the expression of its own entry receptor CD150 (60). In the case of SARS-CoV-2, data from Zheng and colleagues suggested that TLR2 pathway is involved in disease pathogenesis rather than viral control (31).

Although, the exact pathway of the COVID-19 pathogenesis is still unknown, recent data demonstrated that elevated levels of pro-inflammatory cytokines in serum, including CXCL8, is associated with enhanced disease pathogenesis and mortality. Accordingly, inflammatory mediators are promising therapeutic targets to alleviate COVID-19 pathogenesis (20, 61–63). Thus, understanding the molecular determinants responsible for inflammatory cytokine production in the course of SARS-CoV-2 infection could provide future therapeutic targets. Several SARS-CoV-2 components have been described to trigger inflammatory cytokine production including detection of viral RNA by MDA-5 (64), TLR8 (65) and TLR7 (53, 54), activation of ACE-2 by spike (S) protein in epithelial cells (66) and activation of TLR2 by E protein (32). However, the relative contribution of each pathway in immune protection or pathogenesis warrants further studies. It should be noted that the work of Zheng et al showed that unlike the E protein, the S protein does not seem to induce a significant inflammatory reaction (31). This difference underlines the importance of considering the E protein as a therapeutic target. Our findings showed that E protein induced CXCL8 production in TLR2- and NF-κB dependent manner when tested in HEK-TLR2 cell line model. Thus, this model provides an important tool that could be used to screen antagonist compounds which can be used as antiviral drugs. The production of CXCL8, a known neutrophil chemoattractant, is consistent with the reports describing a high circulating neutrophil number and associated injury in the airway and lung tissues in COVID-19 patients (67, 68). Regarding the pathological deleterious effect of CXCL8 in COVID-19 patients, we could consider targeting protein E for therapeutic purposes, either by immunotherapy approaches by administering neutralizing anti-E antibodies to COVID-19 patients in intensive care units (ICU), or by vaccine approach by combining protein E as an immunogen in future vaccine candidates against COVID-19. Indeed, E protein is one of the most conserved in coronaviruses (Lbachir Benmohamed, personnal communication), and could be associated with a crucial function essential for one of the crucial stages of the viral cycle or for the pathogenicity of the virus.

---

**Supplementary Figure S1: Infection of primary human macrophages and VeroE6 cells with NeonGreen SARS-CoV-2 virus:** Primary human macrophages or VeroE6 cell line were infected with NeonGreen SARS-CoV-2 virus (MOI 0.01-1). After 20h of infection-time cells were imaged, inside BSL-3 facility, using EVOS Floïd microscope (Invitrogen). Image show merge of bright field and NeonGreen fluorescence.

## References

1. Hu B, Guo H, Zhou P, Shi ZL. 2021. Characteristics of SARS-CoV-2 and COVID-19. Nature reviews Microbiology 19:141–154.

2. Huang Y, Yang C, Xu XF, Xu W, Liu SW. 2020. Structural and functional properties of SARS-CoV-2 spike protein: potential antivirus drug development for COVID-19. Acta pharmacologica Sinica 41:1141–1149.

3. Letko M, Marzi A, Munster V. 2020. Functional assessment of cell entry and receptor usage for SARS-CoV-2 and other lineage B betacoronaviruses. Nature microbiology 5:562–569.

4. Schoeman D, Fielding BC. 2019. Coronavirus envelope protein: current knowledge. Virology journal 16:69.

5. Kuo L, Hurst KR, Masters PS. 2007. Exceptional flexibility in the sequence requirements for coronavirus small envelope protein function. Journal of virology 81:2249–2262.

6. Baudoux P, Carrat C, Besnardeau L, Charley B, Laude H. 1998. Coronavirus pseudoparticles formed with recombinant M and E proteins induce alpha interferon synthesis by leukocytes. Journal of virology 72:8636–8643.

7. Venkatagopalan P, Daskalova SM, Lopez LA, Dolezal KA, Hogue BG. 2015. Coronavirus envelope (E) protein remains at the site of assembly. Virology 478:75–85.

8. DeDiego ML, Alvarez E, Almazan F, Rejas MT, Lamirande E, Roberts A, Shieh WJ, Zaki SR, Subbarao K, Enjuanes L. 2007. A severe acute respiratory syndrome coronavirus that lacks the E gene is attenuated in vitro and in vivo. Journal of virology 81:1701–1713.

9. Ortego J, Ceriani JE, Patino C, Plana J, Enjuanes L. 2007. Absence of E protein arrests transmissible gastroenteritis coronavirus maturation in the secretory pathway. Virology 368:296–308.

10. Netland J, DeDiego ML, Zhao J, Fett C, Alvarez E, Nieto-Torres JL, Enjuanes L, Perlman S. 2010. Immunization with an attenuated severe acute respiratory syndrome coronavirus deleted in E protein protects against lethal respiratory disease. Virology 399:120–128.

11. Lim KP, Liu DX. 2001. The missing link in coronavirus assembly. Retention of the avian coronavirus infectious bronchitis virus envelope protein in the pre-Golgi compartments and physical interaction between the envelope and membrane proteins. The Journal of biological chemistry 276:17515–17523.

12. Corse E, Machamer CE. 2000. Infectious bronchitis virus E protein is targeted to the Golgi complex and directs release of virus-like particles. Journal of virology 74:4319–4326.

13. Mortola E, Roy P. 2004. Efficient assembly and release of SARS coronavirus-like particles by a heterologous expression system. FEBS letters 576:174–178.

14. DeDiego ML, Nieto-Torres JL, Jimenez-Guardeno JM, Regla-Nava JA, Castano-Rodriguez C, Fernandez-Delgado R, Usera F, Enjuanes L. 2014. Coronavirus virulence genes with main focus on SARS-CoV envelope gene. Virus research 194:124–137.

15. Nieto-Torres JL, Verdia-Baguena C, Jimenez-Guardeno JM, Regla-Nava JA, Castano-Rodriguez C, Fernandez-Delgado R, Torres J, Aguilella VM, Enjuanes L. 2015. Severe acute respiratory syndrome coronavirus E protein transports calcium ions and activates the NLRP3 inflammasome. Virology 485:330–339.

16. Kindler E, Thiel V. 2016. SARS-CoV and IFN: Too Little, Too Late. Cell host & microbe 19:139–141.

17. Kim YM, Shin EC. 2021. Type I and III interferon responses in SARS-CoV-2 infection. Experimental & molecular medicine 53:750–760.

18. Vanderbeke L, Van Mol P, Van Herck Y, De Smet F, Humblet-Baron S, Martinod K, Antoranz A, Arijs I, Boeckx B, Bosisio FM, Casaer M, Dauwe D, De Wever W, Dooms C, Dreesen E, Emmaneel A, Filtjens J, Gouwy M, Gunst J, Hermans G, Jansen S, Lagrou K, Liston A, Lorent N, Meersseman P, Mercier T, Neyts J, Odent J, Panovska D, Penttila PA, Pollet E, Proost P, Qian J, Quintelier K, Raes J, Rex S, Saeys Y, Sprooten J, Tejpar S, Testelmans D, Thevissen K, Van Buyten T, Vandenhaute J, Van Gassen S, Velasquez Pereira LC, Vos R, Weynand B, Wilmer A, Yserbyt J, Garg AD, et al. 2021. Monocyte-driven atypical cytokine storm and aberrant neutrophil activation as key mediators of COVID-19 disease severity. Nature communications 12:4117.

19. Channappanavar R, Perlman S. 2017. Pathogenic human coronavirus infections: causes and consequences of cytokine storm and immunopathology. Seminars in immunopathology 39:529–539.

20. Karki R, Sharma BR, Tuladhar S, Williams EP, Zalduondo L, Samir P, Zheng M, Sundaram B, Banoth B, Malireddi RKS, Schreiner P, Neale G, Vogel P, Webby R, Jonsson CB, Kanneganti TD. 2021. Synergism of TNF-alpha and IFN-gamma Triggers Inflammatory Cell Death, Tissue Damage, and Mortality in SARS-CoV-2 Infection and Cytokine Shock Syndromes. Cell 184:149–168 e117.

21. Jose RJ, Manuel A. 2020. COVID-19 cytokine storm: the interplay between inflammation and coagulation. The Lancet Respiratory medicine 8:e46–e47.

22. Odak I, Barros-Martins J, Bosnjak B, Stahl K, David S, Wiesner O, Busch M, Hoeper MM, Pink I, Welte T, Cornberg M, Stoll M, Goudeva L, Blasczyk R, Ganser A, Prinz I, Forster R, Koenecke C, Schultze-Florey CR. 2020. Reappearance of effector T cells is associated with recovery from COVID-19. EBioMedicine 57:102885.

23. Diao B, Wang C, Tan Y, Chen X, Liu Y, Ning L, Chen L, Li M, Wang G, Yuan Z, Feng Z, Zhang Y, Wu Y, Chen Y. 2020. Reduction and Functional Exhaustion of T Cells in Patients With Coronavirus Disease 2019 (COVID-19). Frontiers in immunology 11:827.

24. He Z, Zhao C, Dong Q, Zhuang H, Song S, Peng G, Dwyer DE. 2005. Effects of severe acute respiratory syndrome (SARS) coronavirus infection on peripheral blood lymphocytes and their subsets. International journal of infectious diseases: IJID: official publication of the International Society for Infectious Diseases 9:323–330.

25. Kieser KJ, Kagan JC. 2017. Multi-receptor detection of individual bacterial products by the innate immune system. Nature reviews Immunology 17:376–390.

26. Broz P, Dixit VM. 2016. Inflammasomes: mechanism of assembly, regulation and signalling. Nature reviews Immunology 16:407–420.

27. Kuriakose T, Man SM, Malireddi RK, Karki R, Kesavardhana S, Place DE, Neale G, Vogel P, Kanneganti TD. 2016. ZBP1/DAI is an innate sensor of influenza virus triggering the NLRP3 inflammasome and programmed cell death pathways. Science immunology 1.

28. Zhang T, Yin C, Boyd DF, Quarato G, Ingram JP, Shubina M, Ragan KB, Ishizuka T, Crawford JC, Tummers B, Rodriguez DA, Xue J, Peri S, Kaiser WJ, Lopez CB, Xu Y, Upton JW, Thomas PG, Green DR, Balachandran S. 2020. Influenza Virus Z-RNAs Induce ZBP1-Mediated Necroptosis. Cell 180:1115–1129 e1113.

29. Bauernfried S, Scherr MJ, Pichlmair A, Duderstadt KE, Hornung V. 2021. Human NLRP1 is a sensor for double-stranded RNA. Science 371.

30. Gringhuis SI, Hertoghs N, Kaptein TM, Zijlstra-Willems EM, Sarrami-Forooshani R, Sprokholt JK, van Teijlingen NH, Kootstra NA, Booiman T, van Dort KA, Ribeiro CM, Drewniak A, Geijtenbeek TB. 2017. HIV-1 blocks the signaling adaptor MAVS to evade antiviral host defense after sensing of abortive HIV-1 RNA by the host helicase DDX3. Nature immunology 18:225–235.

31. Zheng M, Karki R, Williams EP, Yang D, Fitzpatrick E, Vogel P, Jonsson CB, Kanneganti TD. 2021. TLR2 senses the SARS-CoV-2 envelope protein to produce inflammatory cytokines. Nat Immunol 22:829–838.

32. Zheng M, Karki R, Williams EP, Yang D, Fitzpatrick E, Vogel P, Jonsson CB, Kanneganti TD. 2021. TLR2 senses the SARS-CoV-2 envelope protein to produce inflammatory cytokines. Nature immunology 22:829–838.

33. Xie X, Muruato A, Lokugamage KG, Narayanan K, Zhang X, Zou J, Liu J, Schindewolf C, Bopp NE, Aguilar PV, Plante KS, Weaver SC, Makino S, LeDuc JW, Menachery VD, Shi PY. 2020. An Infectious cDNA Clone of SARS-CoV-2. Cell Host Microbe 27:841–848 e843.

34. Planes R, BenMohamed L, Leghmari K, Delobel P, Izopet J, Bahraoui E. 2014. HIV-1 Tat protein induces PD-L1 (B7-H1) expression on dendritic cells through tumor necrosis factor alpha- and toll-like receptor 4-mediated mechanisms. J Virol 88:6672–6689.

35. Bahraoui E, Serrero M, Planes R. 2020. HIV-1 Tat - TLR4/MD2 interaction drives the expression of IDO-1 in monocytes derived dendritic cells through NF-kappaB dependent pathway. Scientific reports 10:8177.

36. Serrero M, Planes R, Bahraoui E. 2017. PKC-delta isoform plays a crucial role in Tat-TLR4 signalling pathway to activate NF-kappaB and CXCL8 production. Scientific reports 7:2384.

37. Baeuerle PA, Baltimore D. 1996. NF-kappa B: ten years after. Cell 87:13–20.

38. Badou A, Bennasser Y, Moreau M, Leclerc C, Benkirane M, Bahraoui E. 2000. Tat protein of human immunodeficiency virus type 1 induces interleukin-10 in human peripheral blood monocytes: implication of protein kinase C-dependent pathway. Journal of virology 74:10551–10562.

39. Li Q, Verma IM. 2002. NF-kappaB regulation in the immune system. Nature reviews Immunology 2:725–734.

40. Vij N, Amoako MO, Mazur S, Zeitlin PL. 2008. CHOP transcription factor mediates IL-8 signaling in cystic fibrosis bronchial epithelial cells. Am J Respir Cell Mol Biol 38:176–184.

41. Hoffmann E, Dittrich-Breiholz O, Holtmann H, Kracht M. 2002. Multiple control of interleukin-8 gene expression. J Leukoc Biol 72:847–855.

42. Mukaida N, Okamoto S, Ishikawa Y, Matsushima K. 1994. Molecular mechanism of interleukin-8 gene expression. J Leukoc Biol 56:554–558.

43. Schwandner R, Dziarski R, Wesche H, Rothe M, Kirschning CJ. 1999. Peptidoglycan- and lipoteichoic acid-induced cell activation is mediated by toll-like receptor 2. The Journal of biological chemistry 274:17406–17409.

44. Yoshimura A, Lien E, Ingalls RR, Tuomanen E, Dziarski R, Golenbock D. 1999. Cutting edge: recognition of Gram-positive bacterial cell wall components by the innate immune system occurs via Toll-like receptor 2. Journal of immunology 163:1–5.

45. Brightbill HD, Libraty DH, Krutzik SR, Yang RB, Belisle JT, Bleharski JR, Maitland M, Norgard MV, Plevy SE, Smale ST, Brennan PJ, Bloom BR, Godowski PJ, Modlin RL. 1999. Host defense mechanisms triggered by microbial lipoproteins through toll-like receptors. Science 285:732–736.

46. Lien E, Sellati TJ, Yoshimura A, Flo TH, Rawadi G, Finberg RW, Carroll JD, Espevik T, Ingalls RR, Radolf JD, Golenbock DT. 1999. Toll-like receptor 2 functions as a pattern recognition receptor for diverse bacterial products. The Journal of biological chemistry 274:33419–33425.

47. Oliveira-Nascimento L, Massari P, Wetzler LM. 2012. The Role of TLR2 in Infection and Immunity. Frontiers in immunology 3:79.

48. Jin MS, Kim SE, Heo JY, Lee ME, Kim HM, Paik SG, Lee H, Lee JO. 2007. Crystal structure of the TLR1-TLR2 heterodimer induced by binding of a tri-acylated lipopeptide. Cell 130:1071–1082.

49. Torres J, Wang J, Parthasarathy K, Liu DX. 2005. The transmembrane oligomers of coronavirus protein E. Biophysical journal 88:1283–1290.

50. Pervushin K, Tan E, Parthasarathy K, Lin X, Jiang FL, Yu D, Vararattanavech A, Soong TW, Liu DX, Torres J. 2009. Structure and inhibition of the SARS coronavirus envelope protein ion channel. PLoS pathogens 5:e1000511.

51. Khanmohammadi S, Rezaei N. 2021. Role of Toll-like receptors in the pathogenesis of COVID-19. Journal of medical virology 93:2735–2739.

52. Lester SN, Li K. 2014. Toll-like receptors in antiviral innate immunity. Journal of molecular biology 426:1246–1264.

53. Cervantes-Barragan L, Zust R, Weber F, Spiegel M, Lang KS, Akira S, Thiel V, Ludewig B. 2007. Control of coronavirus infection through plasmacytoid dendritic-cell-derived type I interferon. Blood 109:1131–1137.

54. Onodi F, Bonnet-Madin L, Meertens L, Karpf L, Poirot J, Zhang SY, Picard C, Puel A, Jouanguy E, Zhang Q, Le Goff J, Molina JM, Delaugerre C, Casanova JL, Amara A, Soumelis V. 2021. SARS-CoV-2 induces human plasmacytoid predendritic cell diversification via UNC93B and IRAK4. The Journal of experimental medicine 218.

55. Kasuga Y, Zhu B, Jang KJ, Yoo JS. 2021. Innate immune sensing of coronavirus and viral evasion strategies. Experimental & molecular medicine 53:723–736.

56. Olejnik J, Hume AJ, Muhlberger E. 2018. Toll-like receptor 4 in acute viral infection: Too much of a good thing. PLoS pathogens 14:e1007390.

57. Planes R, BenMohamed L, Leghmari K, Delobel P, Izopet J, Bahraoui E. 2014. HIV-1 Tat protein induces PD-L1 (B7-H1) expression on dendritic cells through tumor necrosis factor alpha- and toll-like receptor 4-mediated mechanisms. Journal of virology 88:6672–6689.

58. Ben Haij N, Leghmari K, Planes R, Thieblemont N, Bahraoui E. 2013. HIV-1 Tat protein binds to TLR4-MD2 and signals to induce TNF-alpha and IL-10. Retrovirology 10:123.

59. Planes R, Bahraoui E. 2013. HIV-1 Tat protein induces the production of IDO in human monocyte derived-dendritic cells through a direct mechanism: effect on T cells proliferation. PloS one 8:e74551.

60. Bieback K, Lien E, Klagge IM, Avota E, Schneider-Schaulies J, Duprex WP, Wagner H, Kirschning CJ, Ter Meulen V, Schneider-Schaulies S. 2002. Hemagglutinin protein of wild-type measles virus activates toll-like receptor 2 signaling. Journal of virology 76:8729–8736.

61. Jones SA, Hunter CA. 2021. Is IL-6 a key cytokine target for therapy in COVID-19? Nature reviews Immunology 21:337–339.

62. Rubin EJ, Longo DL, Baden LR. 2021. Interleukin-6 Receptor Inhibition in Covid-19 - Cooling the Inflammatory Soup. The New England journal of medicine 384:1564–1565.

63. Mehta P, McAuley DF, Brown M, Sanchez E, Tattersall RS, Manson JJ. 2020. COVID-19: consider cytokine storm syndromes and immunosuppression. Lancet 395:1033–1034.

64. Rebendenne A, Valadao ALC, Tauziet M, Maarifi G, Bonaventure B, McKellar J, Planes R, Nisole S, Arnaud-Arnould M, Moncorge O, Goujon C. 2021. SARS-CoV-2 triggers an MDA-5-dependent interferon response which is unable to control replication in lung epithelial cells. Journal of virology doi:10.1128/JVI.02415-20.

65. Campbell GR, To RK, Hanna J, Spector SA. 2021. SARS-CoV-2, SARS-CoV-1, and HIV-1 derived ssRNA sequences activate the NLRP3 inflammasome in human macrophages through a non-classical pathway. iScience 24:102295.

66. Patra T, Meyer K, Geerling L, Isbell TS, Hoft DF, Brien J, Pinto AK, Ray RB, Ray R. 2020. SARS-CoV-2 spike protein promotes IL-6 trans-signaling by activation of angiotensin II receptor signaling in epithelial cells. PLoS pathogens 16:e1009128.

67. Chen N, Zhou M, Dong X, Qu J, Gong F, Han Y, Qiu Y, Wang J, Liu Y, Wei Y, Xia J, Yu T, Zhang X, Zhang L. 2020. Epidemiological and clinical characteristics of 99 cases of 2019 novel coronavirus pneumonia in Wuhan, China: a descriptive study. Lancet 395:507–513.

68. Qin C, Zhou L, Hu Z, Zhang S, Yang S, Tao Y, Xie C, Ma K, Shang K, Wang W, Tian DS. 2020. Dysregulation of Immune Response in Patients With Coronavirus 2019 (COVID-19) in Wuhan, China. Clinical infectious diseases: an official publication of the Infectious Diseases Society of America 71:762–768.

